# Does the interaction between partnership status and average progesterone level predict women’s preferences for facial masculinity?

**DOI:** 10.1101/376350

**Authors:** Lisa M DeBruine, Amanda C Hahn, Benedict C Jones

## Abstract

Many studies have attempted to identify biological factors that reliably predict individual differences in women’s preferences for masculine male faces. Marcinkowska et al. (2018, Hormones & Behavior) recently reported that women’s (N=102) preferences for facial masculinity were predicted by the interaction between their relationship status (partnered versus unpartnered) and average progesterone level. Because previous findings for between-women differences in masculinity preferences have often not replicated well, we attempted to replicate Marcinkowska et al’s result in an open data set from another recent study that had not tested this hypothesis (Jones et al., 2018 Psychological Science). In this sample of 316 women, we found that facial masculinity preferences were predicted by the interaction between women’s relationship status and average progesterone level, consistent with Marcinkowska et al’s results. Together, these findings suggest that the combined effects of relationship status and average progesterone level may predict facial masculinity preferences relatively reliably.

## Introduction

Masculine facial characteristics in men are hypothesized to signal good genes for immuncompetence and/or dominance, while also signaling antisocial tendencies, such as unwillingness to invest time and other resources in romantic relationships (Little et al., 2011; Penton-Voak et al., 2003). Because characteristics of the perceiver may influence how women resolve this putative trade off between the costs and benefits of choosing a masculine mate (Little et al., 2011; Penton-Voak et al., 2003), many researchers have sought to identify biological factors that might reliably predict individual differences in women’s preferences for facial masculinity (see Zietsch et al., 2015 for a recent review).

Several recent, high-powered studies (e.g., Jones et al., 2018; Marcinkowska et al., 2018; Zietsch et al., 2015) have reported that women’s preferences for facial masculinity do not appear to track within-individual changes in women’s hormone levels in the way that some mate-preference theories have proposed (Gangestad & Simpson, 2000; Penton-Voak et al., 1999; see Jones et al., in press for a review of these recent studies). However, only one of these studies tested if average, rather than daily, hormone levels predicted women’s masculinity preferences (Marcinkowska et al., 2018).

Marcinkowska et al. (2018) reported that the interaction between women’s partnership status and average progesterone levels measured throughout one menstrual cycle predicted facial masculinity preference in a sample of 102 women. Specifically, Marcinkowska et al. (2018) found that average progesterone tended to be negatively correlated with masculinity preferences for women in romantic relationships and tended to be positively correlated with masculinity preferences for women not in romantic relationships.

Findings from studies investigating factors that might predict between-women differences in women’s masculinity preferences have typically replicated poorly (e.g., Zietsch et al., 2015). For example, studies testing whether women using oral contraceptives show stronger preferences for masculine men than do women not using oral contraceptives have variously reported positive, null, and negative results (Cobey et al., 2015; Feinberg et al., 2008; Jones et al., 2018).

In light of the above, the current study analyzed open data from a large study of the hormonal correlates of women’s masculinity preferences (Jones et al., 2018^1^) to establish whether the interaction between partnership status and average progesterone level reported by Marcinkowska et al. (2018) could be replicated in this new, larger data set.

## Methods

### Procedure

Full methods for data collection are reported in Jones et al. (2018). Briefly, 584 young adult women (age: M=21.46 years, SD=3.09 years) judged the attractiveness of ten pairs of male faces (each pair consisting of a masculinized and feminized version of the same face). Images were manipulated by +/-50% of the linear differences in 2D shape between male and female prototype faces using Webmorph (DeBruine, 2017). Participants chose the face in each pair they thought was more attractive. They did this in up to 15 weekly test sessions in which they also provided a saliva sample, reported their partnership status, and reported their hormonal contraceptive use. In each test session, women completed the face-judgment task twice (once assessing men’s attractiveness for a short-term relationship and once assessing men’s attractiveness for a long-term relationship). All face stimuli are publicly available at https://osf.io/9b4y7/.

### Hormone assays

Saliva samples were assayed by Salimetrics UK for progesterone, estradiol, testosterone, and cortisol (see Jones et al., 2018 for details of relevant kits and descriptive statistics).

### Analysis

For comparison with Marcinkowska et al. (2018), only women who reported no use of hormonal contraceptives and did not change their partnership status during the study were included in the final data set (N = 316; unpartnered = 206, partnered = 110). Most of these women (N = 280) completed 5 or more weekly test sessions; 69 women completed 10 test sessions. 227 women were excluded from the initial dataset because they used hormonal contraceptives during the study, 9 women were excluded for missing hormone values, and 48 women were excluded because they changed partnership status during the study.

## Results

Following Marcinkowska et al. (2018), we analyzed our data using a binomial mixed effects model with random intercepts for participants and stimuli.

Because we had participants judge faces in both long-term and short-term contexts in each session, we also included a random intercept for each participant’s session. Random slopes were specified maximally (for context by participant and for the interaction among context, partnership status, and average progesterone by stimulus). The dependent variable was masculinity preference (1 = chose the more masculine face as more attractive, 0 = chose the more feminine face as more attractive) and the predictors were context (effect-coded: short-term = 0.5, long-term = −0.5), partnership status (effect-coded: 0.5 = partnered, −0.5 = unpartnered), and average progesterone for each participant divided by 400 (following Jones et al., 2018, to facilitate model calculations) and centered on the grand mean of the average progesterone values across all participants. The full analysis code and data are available at https://osf.io/q9szc/.

Our model showed significant main effects of context (B = 0.133, SE = 0.046, z = 2.880, p = 0.004, 95% CI = [0.042, 0.223]) and partnership status (B = 0.379, SE = 0.111, z = 3.401, p = 0.001, 95% CI = [0.161, 0.597]). Our model also showed a significant interaction between partnership status and average progesterone (B = −1.447, SE = 0.615, z = −2.353, p = 0.019, 95% CI = [−2.653, −0.242]). This interaction is shown in Figure 1. Separate analyses by partnership status showed that the direction of the relationship between masculinity preference and average progesterone was positive for unpartnered women (B = 0.700, SE = 0.398, z = 1.760, p = 0.078, 95% CI = [-0.080, 1.480]) and negative for partnered women (B = −0.753, SE = 0.505, z = −1.491, p = 0.136, 95% CI = [-1.743, 0.237]).

**Figure 1.**
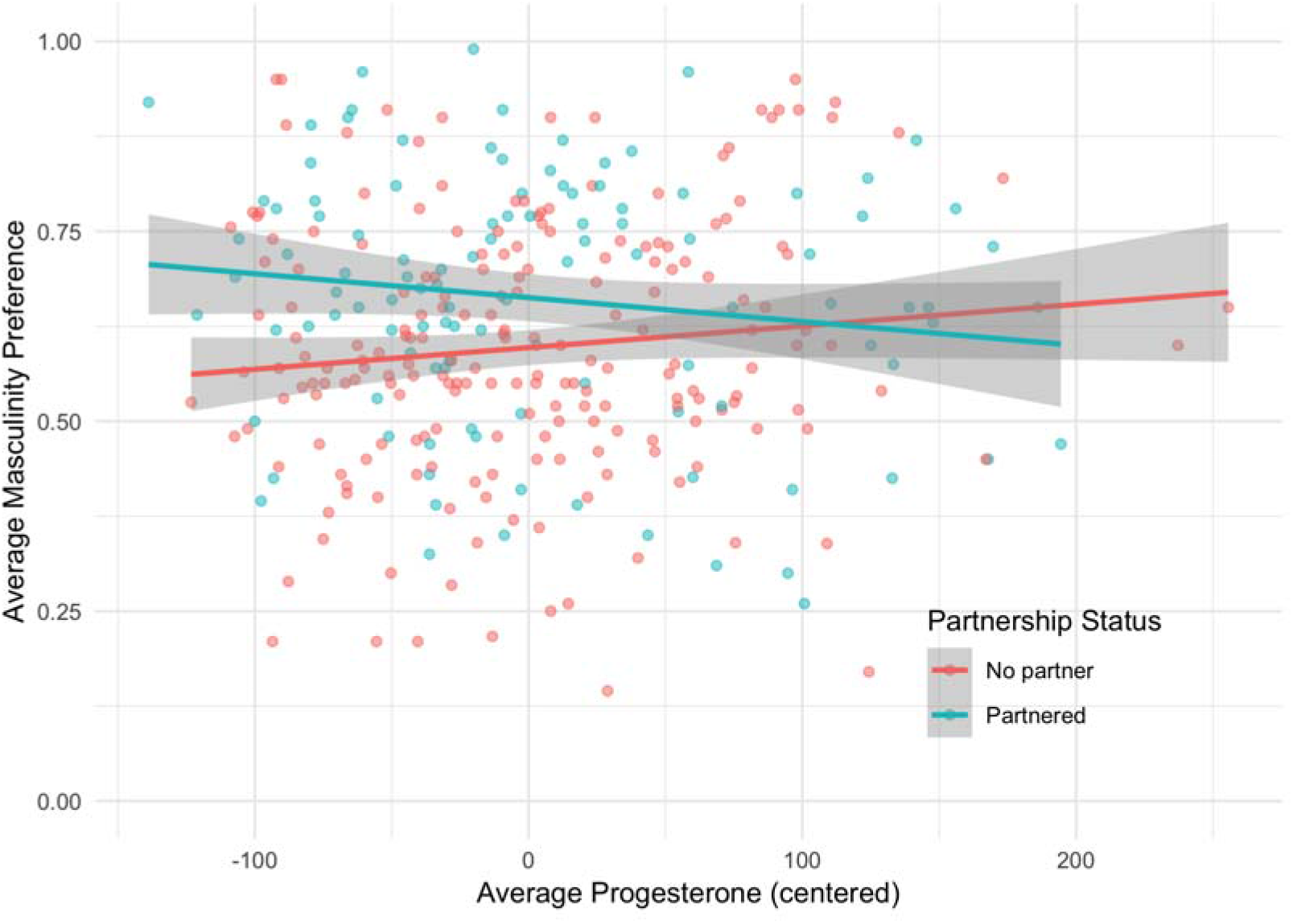
The significant interaction between partnership status and average progesterone.

At the request of the Editor, we repeated this analysis replacing average progesterone with average estradiol, testosterone, or cortisol. Analyses of estradiol showed the same main effects of context (B = 0.130, SE = 0.046, z = 2.845, p = 0.004, 95% CI = [0.041, 0.220]) and partnership status (B = 0.375, SE = 0.112, z = 3.344, p = 0.001, 95% CI = [0.155, 0.595]) described above, but no significant effects involving estradiol (main effect: B = 0.137, SE = 0.268, z = 0.512, p = 0.608, 95% CI = [-0.388, 0.662]; interaction with context: B = −0.291, SE = 0.233, z = −1.249, p = 0.212, 95% CI = [-0.747, 0.166]; interaction with partnership status: B = −0.245, SE = 0.531, z = −0.462, p = 0.644, 95% CI = [-1.286, 0.796]; 3-way interaction with context and partnership status: B = −0.325, SE = 0.440, z = −0.738, p = 0.461, 95% CI = [-1.188, 0.538]).

Analyses of cortisol showed no significant effects including cortisol. Analyses of testosterone showed a significant interaction between average testosterone and context, whereby testosterone tended to be negatively related to masculinity preference in the short-term, but not long-term, condition. The full analysis code, results, and data for these analyses are available at https://osf.io/q9szc/.

## Discussion

Marcinkowska et al. (2018) reported that women’s facial masculinity preferences were predicted by the interaction between partnership status and average progesterone. Here we replicated this finding in a larger sample of women, using open data from Jones et al. (2018). Like Marcinkowska et al. (2018), average progesterone tended to be negatively correlated with masculinity preferences for women in romantic relationships and tended to be positively correlated with masculinity preferences for women not in romantic relationships. Together, our and Marcinkowska et al’s (2018) results suggest that the combined (i.e., interactive) effects of average progesterone level and partnership status predict women’s facial masculinity preferences somewhat reliably. Furthermore, these results highlight the importance of considering possible effects of partnership status when testing for links between hormone levels and women’s masculinity preferences. Given neither Jones et al. (2018) nor Marcinkowska et al. (2018) observed significant within-subject effects of progesterone on masculinity preferences, we suggest it is unlikely that the effects of average progesterone seen here reflect direct (i.e., causal) effects of hormones on mate preferences. Nonetheless, that masculinity preferences appear to be related to average, but not daily, hormone levels is consistent with Havlicek et al. (2015), who proposed that within-women, fertility-linked changes in mating psychology might simply be low-cost functionless byproducts of processes that evolved because of between-women differences in mating psychology (see also Jones et al., in press).

Although we find the same interaction between partnership status and average progesterone reported by Marcinkowska et al. (2018), the simple effects of progesterone for partnered and unpartnered women were not significant in our sample. This difference between our results and Marcinkowska et al’s results could reflect methodological differences. For example, Marcinkowska et al. (2018) took cycle phase into account when calculating average progesterone levels and included only women who showed evidence of ovulation from lutenizing hormone tests in their analyses. Differences in the two samples (rural Poland versus UK university samples) may also contribute to these differences in our results.

In addition to the interaction described above, women preferred masculine men more for short-term relationships than long-term relationships. Partnered women showed stronger preferences for masculine men than did unpartnered women. Both of these results are consistent with previous research reporting effects of relationship context and partnership status on women’s facial masculinity preferences (e.g., Little et al., 2002) and are consistent with the proposal that women show stronger preferences for masculine men in circumstances where investment in the relationship is likely to be less important (e.g., extra-pair relationships, Gangestad & Simpson, 2000). Our null results for average estradiol are also consistent with the null results for estradiol reported by Marcinkowska et al. (2018).

Unexpectedly, a significant interaction between average testosterone and context indicated that women with higher testosterone tended to have weaker masculinity preferences, at least in the short-term rating condition. These results suggest that the relationship between average testosterone and masculinity preferences may be context-dependent. Alternatively, the unexpected interaction between average testosterone and context could be a false positive. Regardless, our results for testosterone contrast with those reported Bobst et al. (2014), who reported that testosterone and masculinity preferences were positively correlated in a sample of 27 women.

In conclusion, here we used open data from a large study testing for within-subject effects of steroid hormones on women’s masculinity preferences (Jones et al., 2018) to replicate the key result from Marcinkowska et al. (2018). Like Marcinkowska et al. (2018), we found that individual differences in women’s facial masculinity preferences were predicted by the interaction between partnership status and average progesterone. Together, these results suggest that the combined effects of these variables on masculinity preferences may be relatively robust.

## Footnotes

1 Jones et al. (2018) tested for within-subject effects of steroid hormones on masculinity preferences, but did not test for between-women effects.

## References

Bobst, C., Sauter, S., Foppa, A., & Lobmaier, J. S. (2014). Early follicular testosterone level predicts preference for masculinity in male faces–but not for women taking hormonal contraception. Psychoneuroendocrinology, 41, 142–150.

Cobey, K. D., Little, A. C., Roberts, S. C. (2015). Hormonal effects on women’s facial masculinity preferences: The influence of pregnancy, post-partum, and hormonal contraceptive use. Biological Psychology, 104, 35–40.

DeBruine, L. M. (2017). Webmorph (Version v0.0.0.9000). Zenodo. http://doi.org/10.5281/zenodo.1073697

Feinberg, D. R., DeBruine, L. M., Jones, B. C., & Little, A. C. (2008). Correlated preferences for men’s facial and vocal masculinity. Evolution & Human Behavior, 29, 233–241.

Gangestad, S. W., & Simpson, J. A. (2000). The evolution of human mating: Trade-offs and strategic pluralism. Behavioral & Brain Sciences, 23, 573–587.

Havliček, J., Cobey, K. D., Barrett, L., Klapilová, K., & Roberts, S. C. (2015). The spandrels of Santa Barbara? A new perspective on the peri-ovulation paradigm. Behavioral Ecology, 26, 1249–1260.

Jones, B. C., Hahn, A. C. & DeBruine, L. M. (in press). Ovulation, sex hormones, and women’s mating psychology. Trends in Cognitive Sciences.

Jones, B. C., Hahn, A. C., Fisher, C. I., Wang, H., Kandrik, M., Han, C., … & O’Shea, K. J. (2018). No compelling evidence that preferences for facial masculinity track changes in women’s hormonal status. Psychological Science, 29, 996–1005.

Little, A. C., Jones, B. C., & DeBruine, L. M. (2011). Facial attractiveness: evolutionary based research. Philosophical Transactions of the Royal Society of London B: Biological Sciences, 366, 1638–1659.

Little, A. C., Jones, B. C., Penton-Voak, I. S., Burt, D. M., & Perrett, D. I. (2002). Partnership status and the temporal context of relationships influence human female preferences for sexual dimorphism in male face shape. Proceedings of the Royal Society of London B: Biological Sciences, 269, 1095–1100.

Marcinkowska, U. M., Kaminski, G., Little, A. C., & Jasienska, G. (2018). Average ovarian hormone levels, rather than daily values and their fluctuations, are related to facial preferences among women. Hormones & Behavior, 102, 114–119.

Penton-Voak, I. S., Perrett, D. I., Castles, D. L., Kobayashi, T., Burt, D. M., Murray, L.K., Minamisawa, R. (1999). Menstrual cycle alters face preference. Nature, 399, 741–742.

Penton-Voak, I. S., Little, A. C., Jones, B. C., Burt, D. M., Tiddeman, B. P., Perrett, D. I. (2003). Female condition influences preferences for sexual dimorphism in faces of male humans (Homo sapiens). Journal of Comparative Psychology, 117, 264–271.

Zietsch, B. P., Lee, A. J., Sherlock, J. M., & Jern, P. (2015). Variation in women’s preferences regarding male facial masculinity is better explained by genetic differences than by previously identified context-dependent effects. Psychological Science, 26, 1440–1448.

